# Protein Cargo Assessment through Residue Alterations

**DOI:** 10.1101/2020.11.20.387738

**Authors:** Ahmed Arslan

**Affiliations:** Department of Anaesthesia, Stanford University School of Medicine, 300 Pasteur drive, Stanford 94035 CA. United States

## Abstract

**Motivation:** The transport of proteins plays a crucial role in the cellular phenotype. Changes in the protein targeting sequence can result in missing protein delivery to the right destination and at the right time, and can disrupt various cellular pathways. Given the importance of single residue change(s) in the protein targeting sequence we developed a missing computational method.

**Results:** By taking into account various protein features like conservation, protein modifications, charge, isoelectric effect and biochemical properties of peptides, the method, *TransSite*, assess the impact of mutations on the protein transportation. We applied this method to big human (cancer proteins) and nonhuman data, and discovered, in both cases, several phenotype related proteins harbour recurring mutations in their targeting sequences.

**Availability:** https://github.com/AhmedArslan/TransSite

**Supplementary information:** Supplementary data are available at *#* online.

## 1 Introduction

The protein transportation from cytoplasm to organelles is a complex process that is tightly regulated by various intrinsic and extrinsic factors. Protein organelle transportation is encoded primarily in sorting region (also known as leading, targeting or signalling sequence) usually present at N-terminal of new peptides and ranges from few to many amino-acids. For example, nuclear targeting signals usually range from 6 to 20 residues whereas endoplasmic reticulum and Golgi bodies targeting protein have signals of 16 to 30 residues long. Similarly, the composition of these sequences varies in their polarity and hydrophobicity. For example, ER sorting proteins have one positive residue followed by a sequence of 6 to 12 amino acids. One such case of protein targeting comes from mitochondrial targeting proteins, the mechanism is complex and modular that is regulated by various components like proteins intrinsic signals as highlighted above, molecular chaperones-based protein sorting and assembly machinery. A change in any component in this pathway can result in the abnormal phenotype. Various diseases have been reported due to the errors in the targeting peptides. Several seminal studies have shown metabolic disease like hyperoxaluria and liver disease (1)(2). Moreover, we recently reported that the mutation carrying TS in mitochondrial inner membrane protein can impact its transportation and result in the fluctuation of metabolite (3). Therefore, knowing the potential impact of amino acid changes that can result in changing the targeting signals in a protein is essential.

We developed a computational method, *TransSite*, that can take into account various crucial aspects of protein targeting sequence (TS) and highlight differences upon the presence of mutations in protein targeting mechanism. *TransSite* is a scoring tool, the score dependent on the change in the (i) amino acid chain charge, (ii) evolutionary conservation (iii) biochemical properties and (iv) protein modifications; upon the presence of the single residue change in the TS. To show its application, we analysed cancer genes and showed that several crucial genes have potential mistargeting due to the presence of mutations in their TS.

## 2 Features

*TranSite* is made available as a python library to the research community. The method searches in the database of proteins with TS for the user provided protein or protein list. Once proteins with TS found for a list of species of interest, the method locate the position(s) of interest in TS containing protein and calculates the following:

### 2.1 Physicochemical properties

Positive charge present at the TS is essential for protein transportation destined for mitochondria. We employ a number of methods to detect the isoelectric point (IP) and charge of TS with and without a mutation. Briefly, IP is pH where the charge of a given protein, or a protein peptide is zero. The IP of a biomolecule depends on the dissociation constant (*pka*) of ionization of charge carrying amino acid residue. The residue charge is intrinsically related to the pH of experimental solution. We employed Henderson-Hasselbalch equation (4) to calculate net charge on a residue, given as:

For positively charged residues:

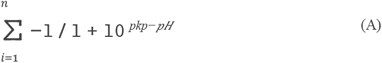

For negatively charged residues:

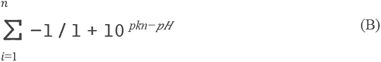

Where *pkp* and *pkn* are dissociation constant of the positively and negatively charged amino acid, respectively. At a given pH, the equation (A) and (B) gives the charge on the protein residues. We utilized the functions of these equations to calculate the charge on the TS in both wild type and altered amino-acid conditions at pre-set pH 7.

### 2.2 Biochemical properties

Diverting the biochemical properties of conserved residues can have a major impact on the protein function and stability. The biochemical properties of can be crucial in the proper function of TS in both protein interaction and cargo to the right destination. We employed various block substitution matrices including Blosum62 to evaluate the differences in the wildtype and alternative amino acids. These methods score the relative probability of biological substitution in a chance event. The inclusion of substitution matrix score can give overall likelihood of whether a residue alteration would impact the cargo activity of TS upon the presence of a mutation.

### 2.3 Conservation

In addition to substitution matrices, we also pre-calculated the rate at which a residue is likely to substitute by other, that depicts the functional importance of a given residue (5). Given the functional estimate of a protein residue, an alternation at a functionally important position could reduce the ability of TS to perform its function(s) properly.

### 2.4 Protein Modifications

Post translational modifications (PTM) play a crucial role in the protein functions especially in protein-protein interactions. Based on the PTM roles, the modified residues present in the TS can play an important role in establishing the interaction with protein transferring chaperones and proper cargo. Changes in PTM state may hamper the TS function.

### 2.5 Implementation

we downloaded the multiple sequence alignment data (6) for ten different species including human (*Homo sapien*), mouse (*Mus musculus*), rat (*Rattus norvegicus*), fruit fly (*Drosophila melanogaster*), frog (*Xenopus laevis*) and yeast (*Saccharomyces cerevisiae*), and precomputed amino-acid residue conservation with rate4Site algorism (5). The pipeline is implemented in a user-friendly python command line, that is available to download from *pip, conda* or *docker*. Two types of inputs are accepted, one is single protein entry with protein names, reference amino acids, mutation position, mutated amino acid (e.g. transsite Lactb M 1 S). Second, is a tab-separated file with entries in the same format as the single protein entry. For the basic use, program takes a protein name, residue position, wildtype residue and alternative residue, and calculate the differences in the aforementioned protein features, to predict the activity of a TS upon the presence of a mutation (**Fig 1**). More negative charge, substitution score and/or conservation score describes the damaging impact of a mutation or vice versa. The final outcome file contains all the score calculations and relative information.

**Figure 1:**
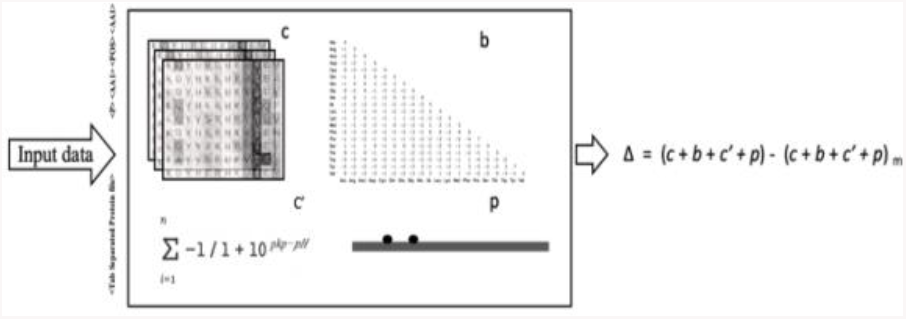
The framework of cargo method. Four different modules are represented, c = conservation, b = block substitution matrix, c’ = charge calculation and p = posttranslational modification. The end product is the difference in the score with (m) and without a mutation.

## 3 Applications

To underscore the applications of *TranSite*, we first focused on the genes reported for human cancers and analysed for the presence of TS, potential change in charge, biochemical and physicochemical properties upon the presence of mutations inside TS. We downloaded (in July 2020) the data from the Catalogue of somatic mutations in cancer (COSMIC) (7). After sorting, and mapping positions to TS, these data consist of ∼11.28 million SNP positions. We found 27 proteins have 39 missense SNPs present in their TS, these proteins, including TFB2M, OTC and PDP1, are reported in various types of cancer including prostate and endometrial cancer (**Table S1**). Another interesting example is of mutation containing protein TS is ACAD6. The higher ACAD9 protein expression reported for its association with the progressive prostate cancer and poorer patient outcomes (8). We found a major amino acid change from Ser to Arg (charge difference= −1), biochemical properties difference (blosum-62 score= −2) of conserved amino acid residue position 2 present inside the TS (Rate4Site score= −0.4779). This interesting mutation can hamper the translocation of ACAD9 to mitochondria which should be validated for its importance as an importance disease marker. In addition to single missense SNP, six other proteins have more than one alternative alleles in their TS. We have recently shown that the presence of multiple mutations in TS ensure the misroute of mitochondrial membrane protein and disrupt normal metabolite level (3) (details in the next section).

A second *TransSite* analysis was performed on the big coding mutational data from 53 inbred mouse strains. We analysed the coding SNP alleles (n=68514) present in 13682 different proteins. A *TransSite* analysis recovered 129 SNP alleles present in the TS of 99 proteins. Out of these cases, with most conserved alleles and biggest changes in the peptide charge, 7 cases of TS SNPs are of interest, which includes Auh, Tmem70, Lactb, Mrpl22, Fxn, Acsm2 and Coq8a proteins (**Table S2**). In case of Lactb, more than one SNP alleles are present inside the TS of mouse Lactb, this protein resides at the inner mitochondrial membrane and known to play a role in organelle metabolism. We recently performed a detailed lab validation of these mutations and shown that the alternative alleles hinder the transportation of the protein, moreover, elevated levels of metabolite succinylcarnitine were also confirmed. The genetic association of LACTB and the metabolite is well established (9) but the contribution of SNPs present inside the TS towards this phenotype was lacking. By taking into the account TS mutated sequence a new mechanism of protein transportation and elevated metabolite levels was uncovered.

Taken together, the importance of alternative alleles carrying TS of genes reported for various human cancers and nonhuman cases suggest previously uncharacterized role of these mutations in the cancer phenotypes. The proteins with mutation(s) in their TS can play a pivotal role in disease(s) and should be studied for further delineate a disease phenotype (**Table S1**).

## Conclusion

We developed an easy to use method to systemically analyse the presence of mutation in the protein targeting sequence and predict functional impact.

## Notes

https://github.com/AhmedArslan/TransSite

### Competing Interest Statement

The authors have declared no competing interest.

