## Supplemental Tables for "Protein Cargo Assessment through Residue Alterations": Table S1.docx

| **Proteins** | **Mutation position** | **∆ Charge** | **∆MW (Da)** | **Substitution score** | **Conservation** | ***TransSite* Prediction** |
| --- | --- | --- | --- | --- | --- | --- |
| Tfb2m | 32 | -1 | 32 | -1 | highly conserved: -1.2067score | Prediction -> likely damaging with major changes |
| Otc | 26 | 1 | 156 | 1 | highly conserved site: 3.1597score | Prediction -> likely naive variation with no consequences |
| Pdp1 | 18 | -1 | 94 | -1 | highly conserved site: -0.2294score | Prediction -> likely damaging with major changes |
| Grsf1 | 72 | 0 | 135 | -2 | variable site: 3.2123 score | Prediction -> possibly damaging |
| Grsf1 | 82 | 1 | 119 | -2 | variable site: 2.7631 score | Prediction -> possibly damaging |
| Coq3 | 19 | -1 | 64 | -2 | highly conserved site: -0.4512score | Prediction -> likely damaging with major changes |
| Nit1 | 26 | 1 | 141 | -1 | variable site: 1.7146 score | Prediction -> possibly damaging |
| Cox5a | 11 | 0 | 133 | -1 | variable site: 2.3973 score | Prediction -> possibly damaging |
| Coq7 | 10 | 0 | 117 | -1 | variable site: 2.2398 score | Prediction -> possibly damaging |
| Mtg1 | 21 | -1 | 60 | -2 | highly conserved site: -0.3849score | Prediction -> likely damaging with major changes |
| Pink1 | 60 | -1 | 14 | -2 | highly conserved site: -0.3911score | Prediction -> likely damaging with major changes |
| Acad9 | 2 | -1 | -12 | -1 | highly conserved site: -0.4779score | Prediction -> likely damaging with major changes |
| Sdhb | 9 | -1 | 156 | -2 | highly conserved site: -0.3462score | Prediction -> likely damaging with major changes |
| Sdhb | 8 | -1 | 69 | -2 | highly conserved site: -0.5405score | Prediction -> likely damaging with major changes |
| Trap1 | 52 | 0 | 137 | -2 | variable site: 2.4551 score | Prediction -> possibly damaging |
| Ppm1k | 19 | 1 | 142 | 0 | variable site: 1.0514 score | Prediction -> possibly damaging |
| Letm1 | 111 | 0 | 31 | -2 | variable site: 1.811 score | Prediction -> possibly damaging |
| Ghitm | 4 | -1 | 99 | -1 | highly conserved site: -0.4120score | Prediction -> likely damaging with major changes |
| Ghitm | 8 | -1 | 85 | -2 | highly conserved site: -0.1939score | Prediction -> likely damaging with major changes |
| Ghitm | 13 | -2 | 70 | -2 | highly conserved site: -0.2469score | Prediction -> likely damaging with major changes |
| Ghitm | 11 | -1 | 159 | -1 | highly conserved site: -0.4407score | Prediction -> likely damaging with major changes |
| Fdx2 | 37 | 0 | 57 | -2 | variable site: 2.3355 score | Prediction -> possibly damaging |
| Star | 22 | -1 | 14 | -2 | highly conserved site: -0.2736score | Prediction -> likely damaging with major changes |
| Star | 43 | 0 | 123 | -1 | variable site: 2.4624 score | Prediction -> possibly damaging |
| Star | 37 | 1 | 141 | 1 | highly conserved site: 2.3593score | Prediction -> likely naive variation with no consequences |
| Hibch | 9 | 0 | 68 | 0 | variable site: 1.0369 score | Prediction -> possibly damaging |
| Spg7 | 34 | 0 | 223 | -2 | variable site: 4.4405 score | Prediction -> possibly damaging |
| Chdh | 28 | 1 | 41 | -3 | variable site: 3.735 score | Prediction -> possibly damaging |
| Pdk4 | 3 | -1 | 101 | -2 | highly conserved site: -0.3550score | Prediction -> likely damaging with major changes |
| Pdk4 | 13 | -1 | 103 | -2 | highly conserved site: -0.3918score | Prediction -> likely damaging with major changes |
| Pdk4 | 28 | -1 | 103 | -2 | highly conserved site: -0.3918score | Prediction -> likely damaging with major changes |
| Pdk4 | 33 | -1 | 119 | -1 | highly conserved site: -0.3918score | Prediction -> likely damaging with major changes |
| Pdk4 | 20 | -1 | 171 | -3 | highly conserved site: -0.3485score | Prediction -> likely damaging with major changes |
| Mecr | 33 | -1 | 39 | -2 | highly conserved site: -0.7149score | Prediction -> likely damaging with major changes |
| Nfu1 | 3 | 1 | 43 | -1 | variable site: 2.0237 score | Prediction -> possibly damaging |
| C1qbp | 21 | 0 | 29 | -2 | variable site: 1.9555 score | Prediction -> possibly damaging |
| C1qbp | 62 | 0 | 17 | -2 | variable site: 2.4647 score | Prediction -> possibly damaging |
| Pdk2 | 24 | -1 | 78 | -1 | highly conserved site: -0.2289score | Prediction -> likely damaging with major changes |
| Clpx | 42 | -1 | 2 | -2 | highly conserved site: -0.1429score | Prediction -> likely damaging with major changes |

**Table S1:** Human somatic mutations reported in various cancer types are analyzed with *TransSite*. 27 proteins have mutation overlap with

targeting sequence (TS), interestingly 6 proteins have recurring mutations (n >1) present in their TS.
