## Supplemental Tables for "Protein Cargo Assessment through Residue Alterations": Table S2.docx

| **Proteins** | **Mutation position** | **∆ Charge** | **∆MW (Da)** | **Substitution score** | **Conservation** | ***TransSite* Prediction** |
| --- | --- | --- | --- | --- | --- | --- |
| Mrpl1 | 9 | 1 | 124 Da | -3 | variable site: 2.033 score | Prediction -> possibly damaging |
| Cars2 | 12 | 0 | 129 Da | -3 | variable site: 2.2136 score | Prediction -> possibly damaging |
| Mrpl55 | 19 | 0 | 37 Da | 0 | variable site: 3.0035 score | Prediction -> possibly damaging |
| Auh | 16 | -1 | -42 Da | -2 | highly conserved site: -0.189 2score | Prediction -> likely damaging with major AA changes |
| Pdpr | 30 | 0 | 167 Da | -1 | variable site: 3.1103 score | Prediction -> possibly damaging |
| Tmem70 | 72 | -1 | 176 Da | -2 | highly conserved site: -0.7170 score | Prediction -> likely damaging with major AA changes |
| Sqor | 6 | 0 | 87 Da | -1 | variable site: 2.0744 score | Prediction -> possibly damaging |
| Lactb | 23 | -1 | 14 Da | -2 | highly conserved site: -0.3218 score | Prediction -> likely damaging with major AA changes |
| Msrb2 | 7 | 0 | 57 Da | -1 | variable site: 2.0653 score | Prediction -> possibly damaging |
| Mrpl22 | 4 | -1 | 85 Da | -1 | highly conserved site: -0.4378 score | Prediction -> likely damaging with major AA changes |
| Ndufs4 | 34 | 0 | 79 Da | 0 | variable site: 2.2515 score | Prediction -> possibly damaging |
| Fxn | 5 | -1 | -42 Da | -2 | highly conserved site: -0.4697 score | Prediction -> likely damaging with major AA changes |
| Dbt | 29 | 0 | 100 Da | -2 | variable site: 2.3989 score | Prediction -> possibly damaging |
| Coq2 | 15 | 0 | 69 Da | 1 | variable site: 0.926 score | Prediction -> possibly damaging |
| Grpel2 | 21 | 0 | 179 Da | -2 | variable site: 3.0597 score | Prediction -> possibly damaging |
| Oxa1l | 42 | 0 | 83 Da | 0 | variable site: 2.1049 score | Prediction -> possibly damaging |
| Acsm2 | 25 | -1 | 122 Da | -1 | highly conserved site: -0.4074 score | Prediction -> likely damaging with major AA changes |
| Sdhc | 4 | 0 | 131 Da | 0 | variable site: 2.0267 score | Prediction -> possibly damaging |
| Naxe | 38 | 0 | 151 Da | -2 | variable site: 2.6125 score | Prediction -> possibly damaging |
| Oxsm | 19 | 0 | 71 Da | -3 | variable site: 3.781 score | Prediction -> possibly damaging |
| Echdc2 | 14 | 0 | 137 Da | -1 | variable site: 2.2032 score | Prediction -> possibly damaging |
| Lars2 | 22 | 1 | 262 Da | -2 | variable site: 2.7186 score | Prediction -> possibly damaging |
| Slc25a3 | 32 | 0 | 81 Da | -1 | variable site: 2.8516 score | Prediction -> possibly damaging |
| Amt | 4 | 0 | 99 Da | -2 | variable site: 3.7287 score | Prediction -> possibly damaging |
| Nfu1 | 15 | 0 | 129 Da | 0 | variable site: 2.2536 score | Prediction -> possibly damaging |
| Cox11 | 17 | 1 | 212 Da | -2 | variable site: 2.295 score | Prediction -> possibly damaging |
| Cox11 | 31 | 0 | 64 Da | -2 | variable site: 2.905 score | Prediction -> possibly damaging |
| Coq8a | 56 | -1 | 14 Da | -2 | highly conserved site: -0.2696score | Prediction -> likely damaging with major AA changes |
| Ndufb2 | 29 | 0 | 167 Da | 0 | variable site: 2.1483 score | Prediction -> possibly damaging |

**Table S2:** Inbred mice protein coding mutations analyzed with *TransSite*. 29 proteins have mutation overlap with targeting sequence (TS), interestingly 7 proteins have recurring mutations (n >1) present in their TS.
